# Cell-intrinsic complement C3 suppresses IFN-β production in macrophages

**DOI:** 10.64898/2026.08.26.746684

**Authors:** Stine Kristensen, Charlotte Årseth, Maria Yurchenko, Liv Ryan, Ida Fjellvær, Kashif Rasheed, Sindre Ullmann, Claudia Kemper, Harald Husebye, Terje Espevik, Trude Helen Flo

**Affiliations:** Department of Clinical and Molecular Medicine, Norwegian University of Science and Technology, Trondheim, Norway; Department of Immunology, Oslo University Hospital, Oslo, Norway; National Heart, Lung, and Blood Institute (NHLBI), National Institutes of Health (NIH), Complement and Inflammation Research Section (CIRS), Bethesda, MD, USA; Clinic of Laboratory Medicine, St. Olavs Hospital, Trondheim, Norway

## Abstract

The cell-intrinsic complement system has emerged as an important orchestrator of a variety of cell-physiological processes, with complement components interacting with intracellular effector systems to regulate cellular responses to pathogens or noxious stimuli. For instance, intracellular C5 signaling through a mitochondrial C5a receptor (C5aR1) controls IL-1β production in human monocytes and macrophages. Here, we investigated whether cell-intrinsic C3 similarly regulates inflammatory responses in macrophages. In LPS-stimulated C3 knockout THP-1-derived macrophages, interferon (IFN)-β production was increased, accompanied by elevated expression of interferon-stimulated genes and enhanced secretion of IFN-induced cytokines and chemokines. C3-deficient cells showed increased phosphorylation of IRF3 at Ser^396^ and a stabilization of the interaction between IRF3 and TBK1, along with enhanced IRF3 dimerization and nuclear translocation. TBK1 phosphorylation was unaffected, indicating that C3 limits IRF3-TBK1 complex formation rather than upstream TBK1 activation. Small-molecule inhibitors of complement factors B and D restored full-length C3 abundance in LPS-stimulated primary human macrophages, consistent with inhibition of the C3 convertase. It also reduced LPS-induced IFN-β production in primary human macrophages and THP-1 cells, suggesting that full-length, uncleaved C3 suppresses IFN-β production. Collectively, these findings identify cell-intrinsic C3 as a suppressor of IFN-β production in human macrophages, highlighting the importance of the cell-intrinsic complement system in fine-tuning inflammatory responses to pathogens.

**One sentence summary:** Cell-intrinsic, full-length C3 suppresses IFN-β production in macrophages by limiting IRF3-TBK1 complex formation, highlighting the role of cell-intrinsic complement in regulating inflammatory responses to pathogens.

## Introduction

The complement system has traditionally been viewed as an effector cascade of liver-derived serum proteins which primarily takes place in the extracellular environment. In response to the detection of pathogens, three different pathways of the complement system may be activated: the classical pathway, the lectin pathway, and the alternative pathway. These pathways all converge on the enzymatic cleavage of the two critical complement components C3 and C5, into their active forms, C3a and C3b, and C5a and C5b. Deposition of C3b on invading microbes and noxious target cells promotes opsonization and phagocytosis via cognate complement receptors on host phagocytes such as macrophages, and drives formation of the C5 convertases, initiating C5b-dependent assembly of the membrane attack complex and lysis of the target cell. Furthermore, stimulation of anaphylatoxin receptors C3a receptor (C3aR) by C3a or C5a receptor 1 (C5aR1) by C5a mediates chemotaxis and the induction of a general inflammatory response [1].

In recent years, a cell-intrinsic and intracellularly active complement system has been discovered in multiple cell types, including T cells, epithelial cells, and macrophages [2–5]. Intracellular complement components and their receptors have been implicated in regulation of multiple cell physiological processes, such as metabolism [6], gene expression [7], and autophagy [8, 9], indicating that the intracellular complement system has a wide variety of non-canonical functions.

We have previously demonstrated that monocytes and macrophages constitutively express C5aR1 on the outer mitochondrial membrane, which is activated by C5a generated by an intracellular C3/C5 convertase [2]. Engagement of mitochondrial C5aR1 triggers a shift in metabolism towards the production of reaction oxygen species and aerobic glycolysis, which in turn favors transcription of the IL1B gene and production of mature IL-1β through activation of the NLR family pyrin domain containing 3 (NLRP3) inflammasome. Although the intracellular presence and cleavage of C3 in human monocytes and macrophages has been established [2–4], it remains unknown whether intracellular C3 plays any role in regulating inflammatory responses in these cells.

Toll-like receptors (TLRs) are a subclass of pattern recognition receptors (PRRs) that crosstalk extensively with the complement system [10] and are rapidly activated upon infection. They respond to an array of pathogen-associated molecular patterns (PAMPs), such as lipopolysaccharide (LPS) from gram-negative bacteria or mycobacterial lipomannans for TLR4 [11]. While some TLRs are restricted to the plasma membrane and others are largely endosomal, TLR4 can be both. Most TLRs signal through the adaptor protein myeloid differentiation primary-response protein 88 (MyD88) to induce nuclear translocation of the transcription factor Nuclear Factor kappa-light-chain-enhancer of activated B cells (NF-κB), and transcription of proinflammatory genes [12]. TLR3 and endosomal TLR4 signal through another adaptor protein; the Toll/IL-1R(TIR)-domain-containing adapter inducing interferon (IFN)-β (TRIF). Following activation and endocytosis of TLR4, the adaptor proteins TRAM and TRIF are recruited to the receptor, which facilitates the activation of the kinases TBK1 and IKKe [13]. In turn, these kinases phosphorylate the transcription factor interferon regulatory factor 3 (IRF3), which then dimerizes and translocates to the nucleus to induce transcription of type I IFNs [14, 15]. Following their release, type I IFNs exert their effects by binding to the interferon-α/β receptor (IFNAR) in an auto- or paracrine fashion [16]. Engagement of IFNAR activates the receptor-associated kinases Janus kinase 1 (JAK1) and tyrosine kinase 2 (TYK2), leading to the downstream phosphorylation and activation of signal transducer and activator of transcription 1 (STAT1) and 2 (STAT2). Activated STAT1/2 proteins dimerize with the transcription factor IRF9, translocate to the nucleus, and then induce the transcription of numerous interferon-stimulated genes (ISGs). This is an integral part of the hosts defense against viruses, bacteria, parasites, and fungi [17]. In line with the reported crosstalk between complement and TLRs, cytosolic C3 has been implicated in NF-κB signaling following stimulation of TLRs in human epithelial cells [18].

Here, we identify a cell-intrinsic role for C3 as a negative regulator of IFN-β production in macrophages. Mechanistically, C3 was found to limit the interaction between IRF3 and TBK1, thereby reducing the phosphorylation of IRF3 at key residues and suppressing its dimerization and nuclear translocation. Furthermore, small molecule inhibitors of the alternative pathway C3 convertase attenuated IFN-β production in LPS-stimulated macrophages, indicating that these effects are mediated by full-length, uncleaved C3. Collectively, our findings reveal a previously unrecognized function of cell-intrinsic C3 in restraining innate immune signaling and underscore the broader importance of noncanonical functions of intracellular complement components in regulating inflammatory responses to pathogens.

## Results

### C3 deficiency increases LPS-induced expression of IFN-β and IFN-β-induced cytokines and chemokines in human and murine macrophages

To investigate the overall effect of C3 knock-out (KO) on transcriptional responses induced by pathogen sensing, we performed bulk RNA sequencing of control and C3 KO THP-1-derived macrophages following stimulation with the TLR4 ligand LPS. Immunoblotting of immunoprecipitated C3 confirmed the absence of C3 protein in KO cells (Fig. S1A). Principal component analysis (PCA) revealed clear separation between control and C3 KO cells under both unstimulated and LPS-stimulated conditions, primarily along principal component 2 (PC2) (Fig. 1A). LPS stimulation induced a shift along PC1 and PC2 in both cell lines. While several genes were differentially expressed in all stimulation conditions, more than a thousand genes were uniquely differentially expressed (both up and down) following LPS stimulation (Fig. 1B). Gene Ontology (GO) enrichment analysis revealed that several pathways related to infections with various pathogens were significantly upregulated in both resting and LPS-stimulated C3 KO cells, including viruses like human papillomavirus, human T-cell leukemia virus, human immunodeficiency virus 1 (HIV-1), and bacterial pathogens like *Salmonella* spp., *Shigella* spp., *Mycobacterium tuberculosis* and *Escherichia coli* (Fig. 1C). Moreover, signaling pathways involved in macrophage activation following pathogen sensing, including PI3K-Akt and MAPK signaling [19, 20], were significantly upregulated in the C3 KO cells. Downregulated signaling pathways included pathways related to cancer, cytokine-cytokine receptor-interaction, NF-κB- and TNF signaling (Fig. S1B).

**Figure 1:**
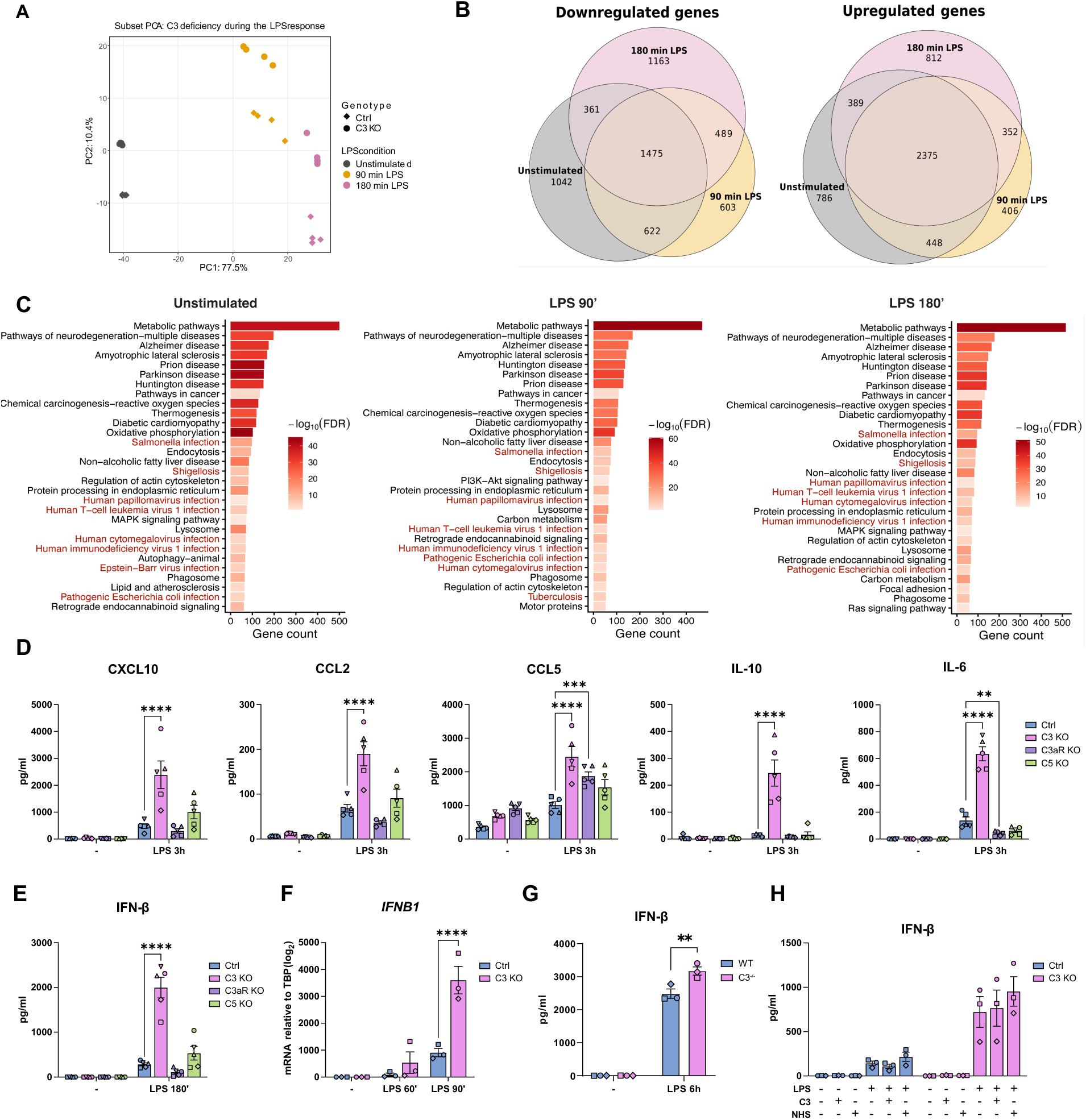
C3 deficiency increases LPS-induced expression of IFN-β and IFN-β-induced cytokines and chemokines in human and murine macrophages. A) Principial component analysis (PCA) of RNA-sequencing data for control and C3 KO THP-1-derived macrophages stimulated as indicated. n = 5. B) Venn diagram of the number of up- and downregulated genes in C3 KO compared to control cells. n = 5. C) KEGG pathway analysis of upregulated pathways in C3 KO cells compared to control cells, showing the top 30 upregulated pathways (adjusted p-value < 0.05, |log2FC| > 2). Pathways related to pathogenic infection highlighted in red. n = 5. D) Quantification of CXCL10, CCL2, CCL5, IL-10, and IL-6 by multiplex immunoassay of supernatants from THP-1 control, C3 KO, C3aR KO and C5 KO cells following stimulation with LPS for 3 h. n = 5. E) Quantification of IFN-β levels by ELISA of supernatants from control-, C3 KO-, C3aR KO- and C5 KO THP-1 cells following stimulation with LPS for 3 h. n = 5. F) Quantification of *IFNB1* mRNA levels by RT-qPCR in control and C3 KO THP-1 cells stimulated with LPS as indicated. n = 3. G) Quantification of IFN-β levels by ELISA of supernatants from control- and C3 KO THP-1 cells, pre-treated for 1 h with serum-purified C3 or 10% normal human serum (NHS) prior to LPS stimulation for 3 h. n = 3. H) Quantification of IFN-β levels from multiplex immunoassay of supernatants from BMDMs made from WT and C3 KO (C3^-/-^) C57BL/6 mice, measured after 6 h of stimulation with LPS. n = 3. Data are expressed as mean ± SEM, with n denoting biological replicates. Statistical significance was determined using two-way ANOVA and Dunnett’s multiple comparisons test (D, E), Šídák’s multiple comparisons test (F, G), or Tukey’s multiple comparisons test (H). * = p < 0.05, ** = p < 0.01, *** = p < 0.001, **** = p < 0.0001.

To further assess the role of C3 in the regulation of inflammatory responses, a multiplex cytokine assay was used to quantify the release of key cytokines and chemokines following LPS stimulation. C3aR KO and C5 KO cells were also included to determine whether an effect on cytokine production could be caused by downstream signaling by C3a or C5 cleavage products. The release of several cytokines and chemokines was significantly increased in the C3 KOs compared to the control cells, including CXCL10, CCL2, CCL5, IL-10, and IL-6 (Fig. 1D). In contrast, except for increased CCL5 and reduced IL-6 in the C3aR KO cells, no significant changes were seen in the C3aR and C5 KO cells. Other cytokines tested were mostly unaffected by C3 KO, excluding a general effect on cytokine secretion in these cells (Fig. S1C). As all the upregulated cytokines in Fig. 1D can be induced or enhanced by IFN-β [21–25], we next measured LPS-induced IFN-β in the C3 KO and control THP-1-derived macrophages by ELISA. IFN-β release was significantly increased in C3 KOs compared to control cells after 3 h of LPS stimulation, but not in C3aR or C5 KO cells (Fig. 1E). Additionally, *IFNB1* mRNA levels were significantly elevated in the C3 KO cells after 90 min of LPS stimulation (Fig. 1F). We next isolated bone marrow-derived macrophages (BMDMs) from wild type (WT) and C3 KO C57BL/6J (C3^-/-^) mice. After 6 h of LPS stimulation, the release of IFN-β, CCL5, CXCL1 and IL-10 was significantly higher in the C3^-/-^ BMDMs compared to the WT (Fig. 1G and Fig. S1D), suggesting that the regulatory role of C3 in IFN-β production is shared across species.

Finally, we investigated whether exogenous C3 could rescue the effects of C3 deficiency by adding serum-purified C3 or normal human serum (NHS) to control and C3 KO cells prior to LPS stimulation. Addition of serum-purified C3 or NHS did not significantly change IFN-β release in either control or C3 KO cells (Fig. 1H), consistent with cell-intrinsic rather than extracellularly sourced C3 acting to suppress IFN-β production. Together, these results indicate that cell-intrinsic C3 suppresses IFN-β production in both human and murine macrophages, likely independent of C3aR signaling and C3-dependent cleavage of C5.

### Signaling through IFNAR and transcription of ISGs is increased in C3-deficient macrophages

Engagement of IFNAR by type I IFNs triggers the phosphorylation and activation of STAT1 and STAT2 by JAK1 and TYK2 kinases, leading to transcription of ISGs [17]. Consistent with increased IFN-β secretion, LPS-induced phosphorylation of STAT1 at Tyr^701^ was significantly increased in C3 KO compared with control THP-1-derived macrophages (Fig. 2A). Furthermore, our transcriptomic data revealed that several ISGs were significantly up- or downregulated in the C3 KO compared to control cells, even at resting conditions (Fig. 2B). After stimulation with LPS, the percentage of upregulated ISGs in the C3 KO increased (Fig. 2C). Among these were key antiviral effectors such as ISG15 and genes in the Interferon-Induced protein with Tetratricopeptide repeats (IFIT) family, the large GTPases GBP1 and MX2, and chemokines and cytokines CXCL9, CXCL10, CXCL11, and IL6 [17] (Fig. 2D). The downregulated ISGs were related to apoptosis ([26, 27], immunosuppressive environments [28], IFN inhibition [29, 30], oncogenes and tumor-suppressors [31, 32], and T cell differentiation and activation [33–35]. To test whether the elevated cytokine release in C3 KO cells depends on autocrine/paracrine type I IFN signaling, cells were pretreated with a blocking antibody against IFNAR prior to LPS stimulation. IFNAR blockade reduced CXCL10 release in C3 KO cells to levels comparable to those in the control cells (Fig. 2E). Collectively, these results indicate that the elevated production of IFN-regulated cytokines in C3 KO cells is mediated by enhanced IFN-β signaling through IFNAR.

**Figure 2:**
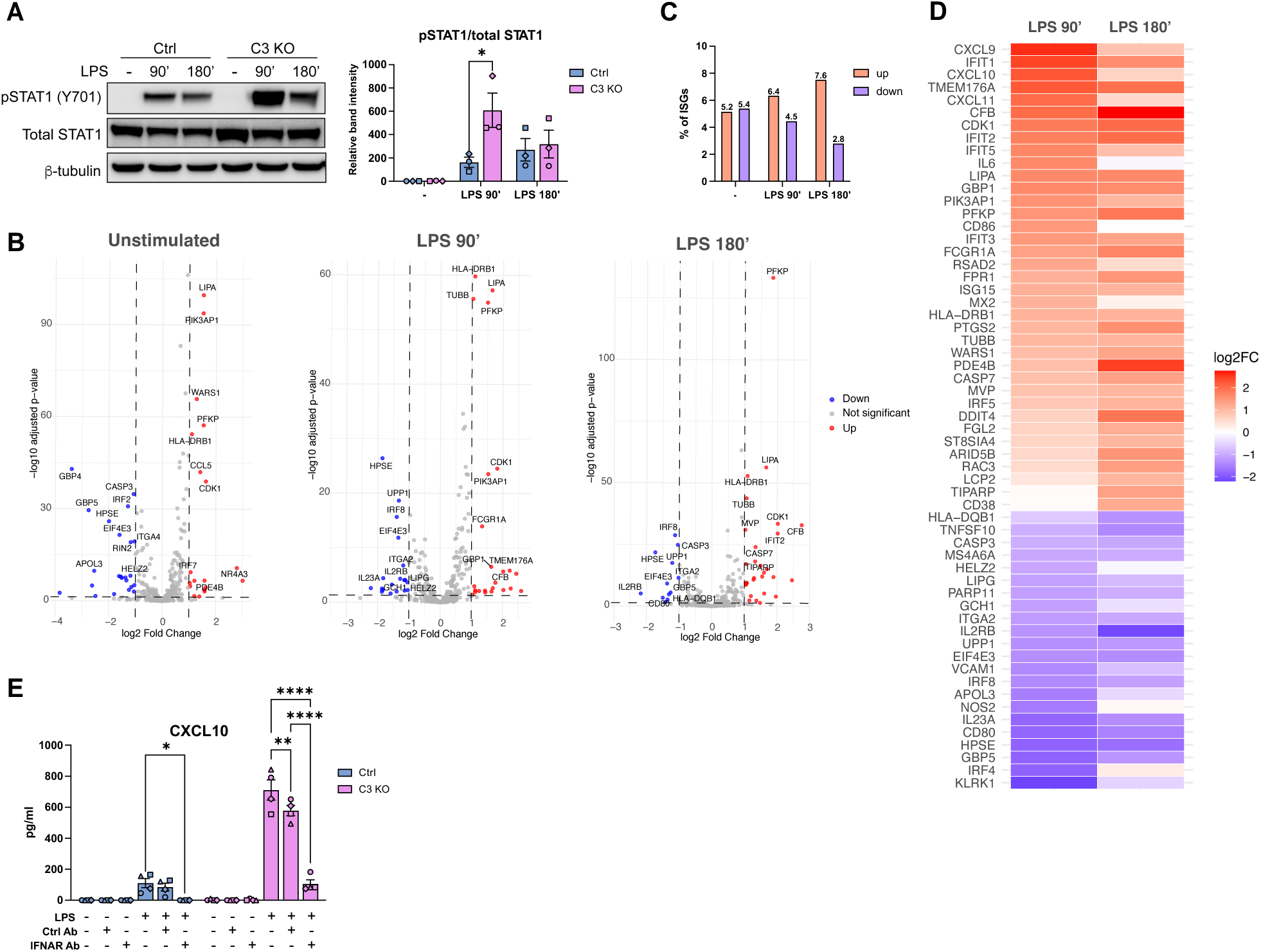
Signaling through IFNAR and transcription of ISGs is increased in C3-deficient macrophages. A) Immunoblot of phospho-STAT1 (Y701), total STAT1 and b-tubulin in lysate from THP-1 control, C3 KO, C3aR KO, and C5 KO cells. Quantifications are based on densitometric analysis and normalised to the corresponding b-tubulin band intensity. n = 3. B) Volcano plot showing log2FC of all ISGs which were significantly altered in the C3 KO cells when compared to control cells (adjusted p-value < 0.05, |log2FC| ≥ 1). C) Quantification of the differentially expressed ISGs from D). D) Heatmap of all ISGs which were differentially expressed after either 90 or 180 min of stimulation with LPS in D), sorted by decreasing log2FC for LPS 90 min. E) Quantification of CXCL10 levels in supernatants by ELISA from control and C3 KO THP-1 cells pre-treated for 30 min with control or IFNAR blocking antibody prior to stimulation with LPS for 3 h. n = 4. B-D from the same RNA sequencing experiments, n = 5. n denotes biological replicates. In A and E, data are expressed as mean ± SEM. n denotes biological replicates. Statistical significance was determined using two-way ANOVA and Dunnett’s multiple comparisons test (A), or Tukey’s multiple comparisons test (E). * = p < 0.05, ** = p < 0.01, *** = p < 0.001, **** = p < 0.0001

### Inhibitors of the alternative C3 convertase decrease the production of IFN-β in THP-1 cells and primary hMDMs

The canonical functions of C3 are induced by the cleavage of C3 into C3a and C3b by a C3 convertase [1]. Human monocytes contain an intracellular alternative pathway C3/C5 convertase [2], and the canonical formation of this convertase depends on the factor D (FD)-mediated cleavage of factor B (FB) into the fragments Ba and Bb [36]. While our results in Fig. 1 indicated that the C3a/C3aR axis is not involved in C3-mediated regulation of IFN-β production, other cleavage fragments, such as C3b, could be responsible. To assess the role of C3 cleavage, primary human monocyte-derived macrophages (hMDMs) were pre-treated with a cell-permeable inhibitor of FB [37] prior to stimulation with LPS. Inhibition of FB caused a significant decrease in the expression of *IFNB1* mRNA in hMDMs (Fig. 3A and Fig. S2A) while IFN-β protein levels in the supernatant displayed a similar trend (Fig. S2B). Furthermore, LPS stimulation decreased the levels of full-length C3 in lysates from untreated cells, but not from FB inhibitor-treated cells, indicating that FB inhibition reduced LPS-induced C3 cleavage (Fig. 3B). Inhibition of extracellular C3 cleavage by a FB-blocking antibody [38], which is unlikely to cross the cell membrane, did not significantly alter the secretion of IFN-β in WT THP-1 cells (Fig. 3C). We next pretreated WT THP-1 cells and primary hMDMs with a small-molecule inhibitor of FD serine protease activity before stimulation with LPS. FD inhibition significantly reduced IFN-β secretion in THP-1 cells (Fig. 3D) and decreased *IFNB1* mRNA expression in hMDMs (Fig. 3E and Fig. S2C). In hMDMs, FD inhibition also reduced LPS-induced cleavage of C3 (Fig. 3F). Together with the factor B inhibitor data, these results show that inhibiting components of the alternative pathway reduces LPS-induced C3 cleavage and IFN-β production in both THP-1-derived and primary human macrophages, suggesting that full-length C3 suppresses IFN-β production.

**Figure 3:**
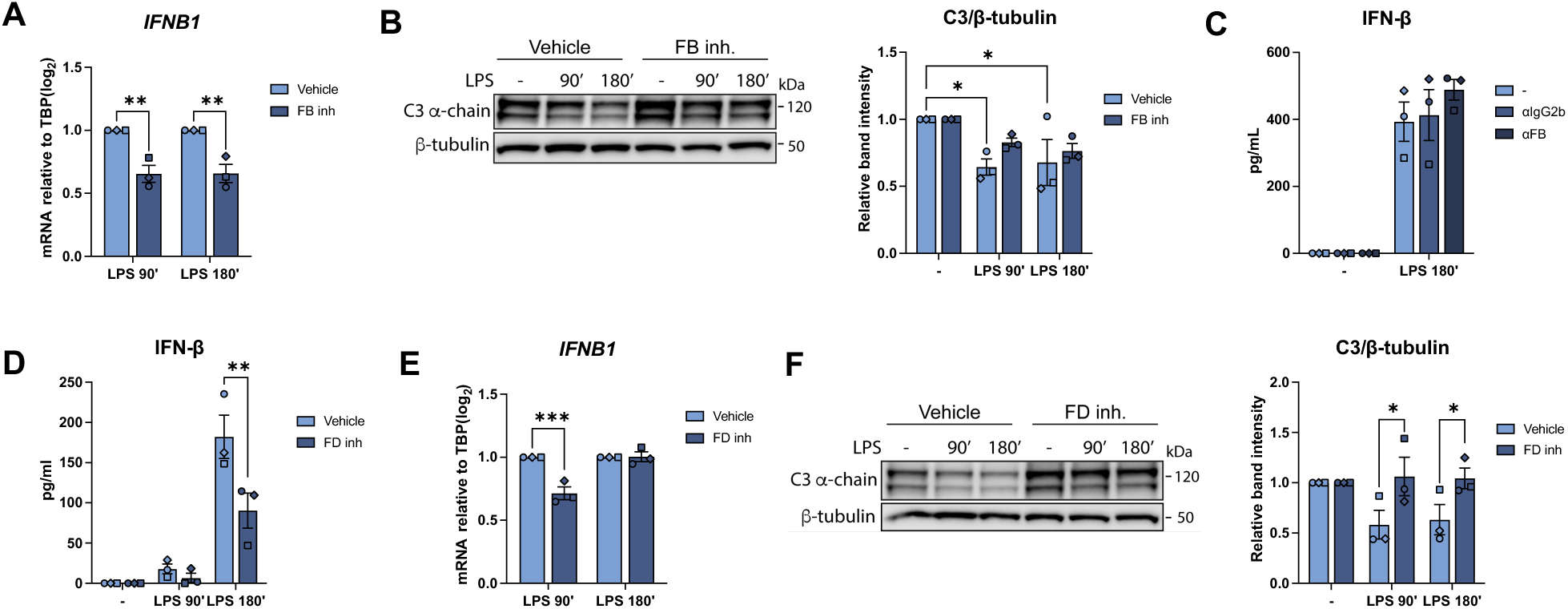
Inhibitors of the alternative C3 convertase decrease the production of IFN-β in THP-1 cells and primary hMDMs. A) Quantification of *IFNB1* mRNA by RT-qPCR of primary hMDMs which have been pre-treated with a cell-permeable FB inhibitor for 1 h prior to LPS stimulation for 90 or 180 min. Inhibitor-treated samples were normalized to their respective vehicle controls. N = 3. B) Immunoblot of total C3 levels in lysate from primary hMDMs from A). Quantifications of C3 α-chain intensities were normalised using b-tubulin intensities. n = 3. C) Quantification of IFN-β protein levels by ELISA of supernatants from THP-1 WT pre-treated with a control antibody or antibody against FB for 1h, followed by 180 min of LPS stimulation. n = 3. D) Quantification of IFN-β secretion by ELISA of supernatants from THP-1 WT pre-treated with FD inhibitor and stimulated with LPS as indicated. n = 3. E) Quantification of *IFNB1* mRNA by RT-qPCR of primary hMDMs pre-treated with FD inhibitor for 1 h prior to LPS stimulation as indicated. Inhibitor-treated samples were normalized to their respective vehicle controls. n = 3. F) Immunoblot of total C3 levels in lysate from primary hMDMs from E). Quantifications of C3 α-chain intensities were normalised using b-tubulin intensities. n = 3. Quantifications of all immunoblots are based on densiometric analysis. Data are expressed as mean ± SEM, n denotes biological replicates. Statistical significance was determined using two-way ANOVA and Šídák’s multiple comparisons test (A, C, D, E) or Tukey’s multiple comparisons test (B, F). * = p < 0.05, ** = p < 0.01, *** = p < 0.001, **** = p < 0.0001.

### C3 deficiency enhances LPS-induced phosphorylation of IRF3 Ser^396^ and subsequent dimerization and nuclear translocation of IRF3

To elucidate the molecular mechanisms by which C3 attenuates IFN-β release, we next assessed how C3 deficiency affects the different steps in the TLR4 signaling pathway regulating IFN-β expression. The enhanced production of IFN-β and IFN-β-induced cytokines and chemokines in C3 KO THP-1-derived macrophages could not be explained by increased expression of TLR4, since *TLR4* mRNA expression was reduced in these cells (Fig. S3A). Furthermore, treatment of the C3 KO cells with the TBK1-IKKe inhibitor MRT67307 significantly reduced IFN-β secretion to levels below the limit of detection, indicating that the increased IFN-β production stems from canonical signaling through TBK1 (Fig. 4A).

**Figure 4:**
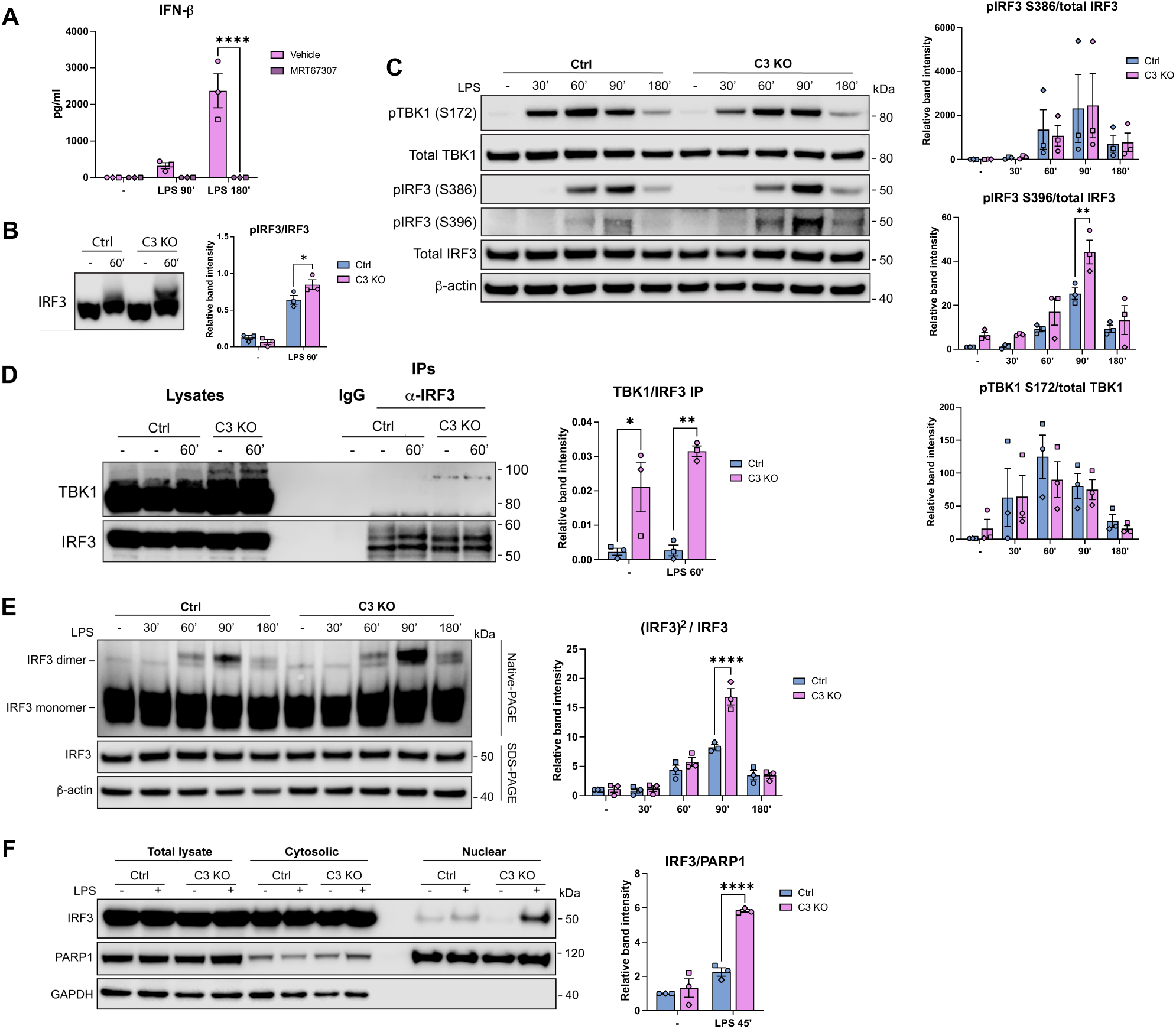
C3 deficiency enhances LPS-induced phosphorylation of IRF3 Ser^396^ and subsequent dimerization and nuclear translocation of IRF3. A) Quantification by ELISA of IFN-β levels in supernatants from C3 KO THP-1 cells pre-treated with MRT67307 for 1 h prior to stimulation with LPS for 3 h. n = 3. B) SDS-PAGE immunoblots with Phos-Tag gels showing total phosphorylation IRF3 in control and C3 KO THP-1-derived macrophages after stimulation with LPS for 60 min. Quantifications are of phosphorylated bands relative to non-phosphorylated bands. n = 3. C) Immunoblots of control and C3 KO THP-1 cells after stimulation with LPS as indicated. Quantifications of phospho-sites are normalized to total protein and b-actin loading control. n = 3. D) Co-IP of endogenous IRF3 and TBK1 from control and C3 KO THP-1-derived macrophages after 60 min of LPS stimulation. Quantifications of the TBK1 bands in IPs were normalised to the intensity of the IRF3-IP bands. n = 3. E) Native PAGE immunoblots of IRF3 dimer formation in lysates from control and C3 KO THP-1 cells stimulated with LPS as indicated. SDS-PAGE of IRF3 and b-tubulin also shown from the same lysates. For quantification, IRF3 dimers were normalised to IRF3 monomers. n = 3. F) Immunoblot of total lysate, cytosolic fractions and nuclear fractions from control and C3 KO THP-1 cells stimulated with LPS for 45 min. n = 3. Quantifications of IRF3 levels in the nucleus were done using nuclear PARP1 for normalization. n = 3. Data are expressed as mean ± SEM, with n denoting biological replicates. Quantifications of all immunoblots are based on densiometric analysis. Statistical significance was determined using either two-way ANOVA and Šídák’s multiple comparisons test (A, C-F) or Unpaired t test with Welch’s correction (B). * = p < 0.05, ** = p < 0.01, *** = p < 0.001, **** = p < 0.0001.

Next, to evaluate the activation of IRF3, the overall phosphorylation profile of IRF3 in the THP-1 C3 KO and the control cells was analyzed by SDS-PAGE with Phos-Tag gels. These gels contain Phos-Tag acrylamide, a phosphate-binding tag which allows for separation of proteins based on their phosphorylation state [39]. Phos-Tag SDS-PAGE demonstrated an upward shift in IRF3 following LPS stimulation, which was significantly stronger in C3 KO cells compared to control cells (Fig. 4B), indicating increased phosphorylation of IRF3 in the absence of C3. The C-terminal loop of IRF3 contains two different phosphorylation sites, with the 2S site consisting of residues Ser^385^ and Ser^386^, and the 5ST consisting of residues Ser^396^, Ser^398^, Ser^402^, Thr^404^, and Ser^405^ [40]. To identify which sites were responsible for the increased IRF3 phosphorylation observed in C3 KO cells, we used phospho-site-specific antibodies against key residues from each site; Ser^386^ and Ser^396^. Phosphorylation of Ser^396^, but not Ser^386^, was significantly increased in C3 KOs after stimulation with LPS for 90 min (Fig. 4C). Activation of TBK1, measured as phosphorylation of Ser^172^ [41], was not significantly altered in the C3 KO cells, demonstrating that the increased IRF3 phosphorylation is not due to hyperactivation of TBK1. Moreover, neither transcription of IRF3 (Fig. S3B), nor total protein levels of IRF3 or TBK1 were altered in the C3 KO cells (Fig. S3C). We next asked whether C3 instead affects the physical interaction between IRF3 and TBK1, and immunoprecipitated IRF3 from lysates of THP-1 control and C3 KO cells. TBK1 co-precipitated with IRF3 in lysates from both resting and LPS-stimulated C3 KO cells, but not in lysates from control cells (Fig. 4D), indicating that the interaction between IRF3 and TBK1 is destabilized by C3. Furthermore, C3 co-precipitated with IRF3, but not TBK1, in lysates from WT cells expressing EGFP-tagged C3 (Fig. S3D-E). These results suggest that a physical interaction between C3 and IRF3 may in turn limit the interaction between IRF3 and TBK1.

Phosphorylation of IRF3 at Ser^386^ and Ser^396^ induces its dimerization and subsequent nuclear translocation [15]. Native PAGE revealed that the ratio of IRF3 dimers to IRF3 monomers was significantly increased in C3 KO cells compared to control cells after 90 min of LPS stimulation (Fig. 4E). We assessed IRF3 nuclear translocation by immunoblotting cytosolic and nuclear fractions of LPS-stimulated control and C3 KO cells. Nuclear fractions of C3 KO cells contained significantly higher levels of IRF3 than the control cells after 45 min of LPS stimulation (Fig. 4F), indicating enhanced nuclear translocation of IRF3 in the absence of C3. In contrast, nuclear translocation of transcription factors IRF1, IRF7, and NF-κB p65 was not significantly altered in the C3 KO cells (Fig. S3F). Collectively, these results suggest that C3 suppresses IFN-β expression by limiting complex formation between IRF3 and TBK1, thereby reducing TBK1-dependent phosphorylation of IRF3 Ser^396^ and subsequent IRF3 dimerization and nuclear translocation.

### C3 deficiency increases IFN-β release upon infection with *E. coli* and *Mycobacterium tuberculosis*

We next asked whether C3-mediated suppression of IFN-β extends to infection with live bacteria like *E. coli*, the pathogen from which the LPS used in this study is derived. After 4 h of infection with the non-virulent *E. coli* strain DH5α, IFN-β release was significantly increased in C3 KO compared to control THP-1-derived macrophages (Fig. 5A), consistent with our observations after LPS stimulation. This was not due to differences in cellular survival, as lytic cell death measured by release of lactate dehydrogenase (LDH), was not altered in the C3 KO cells (Fig. 5B). Having confirmed that C3 restrains IFN-β production during infection with a non-virulent bacterium, we next sought to test whether this effect extends to a clinically relevant, virulent pathogen. Guided by our transcriptomic data, in which the KEGG pathway “Tuberculosis” was significantly upregulated in C3 KO cells (Fig. 1), we selected *Mycobacterium tuberculosis* (*Mtb*) for further infection experiments. *Mtb* can induce type I IFNs through several different surface- and cytosolic PRRs, including TLR4, and type I IFN signaling is thought to contribute to tuberculosis (TB) pathology [42, 43]. Upon infection with auxotrophic *Mtb* H37Rv mc²6206, IFN-β secretion in C3 KO cells trended upward at all timepoints, with a significant increase at 18 h post-infection (Fig. 5C). Lytic cell death was not affected by C3 KO in *Mtb* infection (Fig. 5D). Moreover, neither *Mtb* uptake (Fig. 5E), nor intracellular survival (Fig. 5F) were altered in the C3 KO cells compared to the controls. Key findings were confirmed in BMDMs, where secretion of IFN-β and IL-6 was significantly increased 6 h post-infection in *Mtb*-infected C3^-/-^ BMDMs (Fig. 5G, Fig S4). Collectively, these findings demonstrate that the role of C3 in attenuating IFN-β production extends beyond the TLR4-ligand LPS to more clinically relevant infections with live bacteria

**Figure 5:**
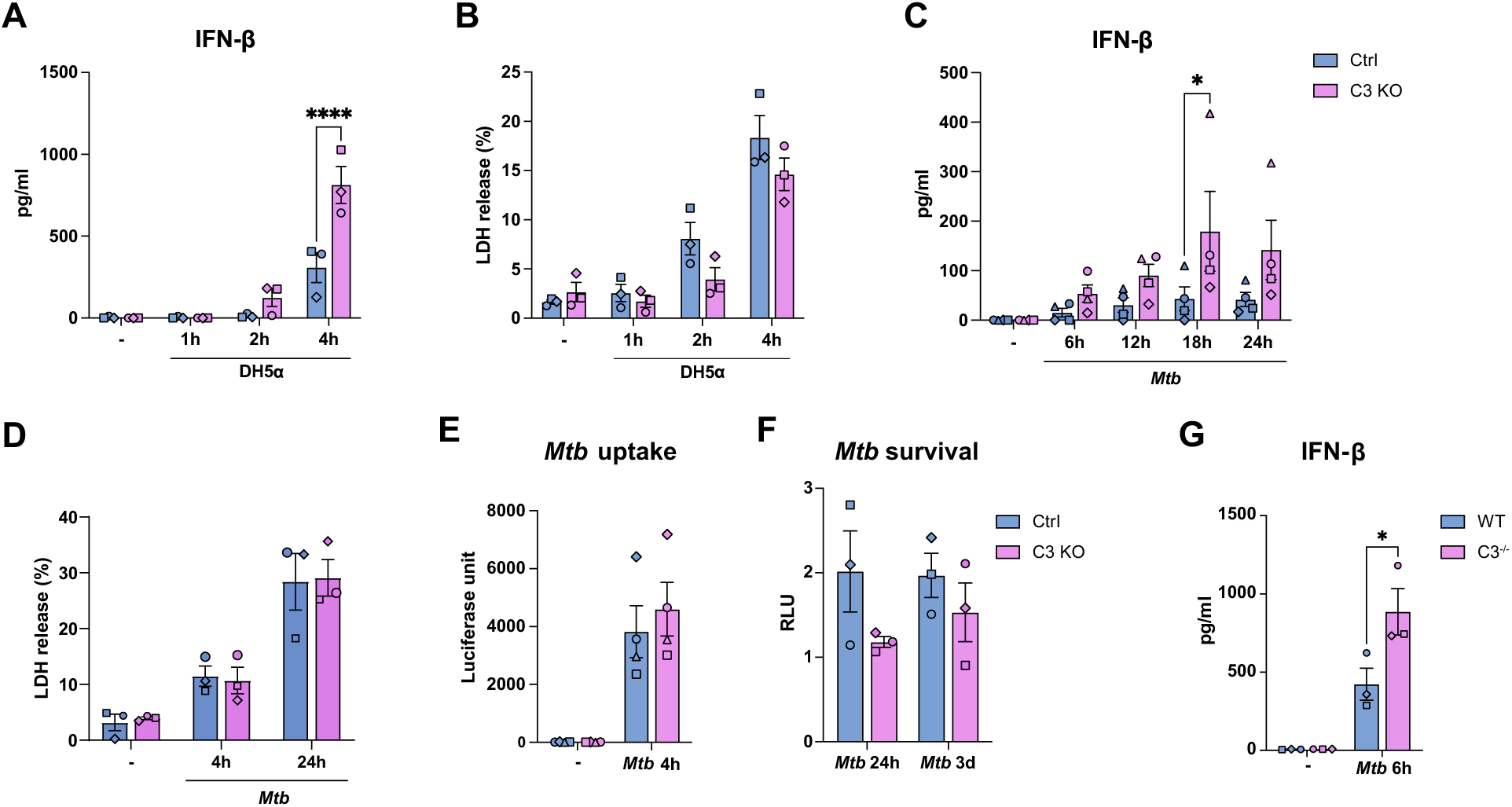
C3 deficiency increases IFN-β release upon infection with *E. coli* and M. tuberculosis. A) Quantification of protein levels of IFN-β in supernatants from control and C3 KO THP-1-derived macrophages after infection with *E. coli* for 1, 2, and 4 h. n = 3. B) Cell death measured by LDH release in the supernatants from (A). n = 3. C) Quantification of IFN-β release by ELISA from supernatants from control and C3 KO THP-1-derived macrophages after infection with auxotroph *Mycobacterium tuberculosis* (*Mtb*) for 6, 12, 18 or 24 h. n = 4 D) Cell death measured by LDH release in control and C3 KO THP-1 cells after stimulation with auxotroph *Mtb* for 4 and 24 h. n = 3. E) Uptake of *Mtb* at 4 h, measured by LU. n = 4. F) Intracellular survival of auxotroph *Mtb* for 24 h and 3 days in control- and C3 KO THP-1 cells, normalized to cellular uptake of *Mtb* at 4 h and measured as relative luciferase light units (RLU). G) Quantification of IFN-β secretion in supernatants from BMDMs from WT and C3^-/-^ C57BL/6 mice after infection with auxotroph *Mtb* for 6 h, measured by multiplex immunoassay n = 3. Data are expressed as mean ± SEM, n denotes biological replicates. Statistical significance was determined using two-way ANOVA and Šídák’s multiple comparisons test. * = p < 0.05, ** = p < 0.01, *** = p < 0.001, **** = p < 0.0001.

## Discussion

The discovery of cell-intrinsic complement as a critical regulator of basic cell physiology [1, 4, 7, 8], cell death [5, 44], and infection [9, 18, 45, 46] has caused a paradigm shift in complement biology, expanding our understanding of complement beyond its classical role as the serum-effective arm of innate immunity.

Here, we show that full-length C3 suppresses the production of IFN-β by interfering with IRF3 signaling downstream of TLR4 activation in macrophages. Complement and TLRs are both crucial components of the innate immune system, and molecular interplay between these signaling pathways may have evolved to provide increased specificity and sensitivity to our first-line defense against infection. As such, complement regulates TLR signaling pathways in multiple ways [10]. For instance, C5a has been shown to attenuate TLR4-induced expression of cytokines in the IL-12 family [47, 48], and simultaneous addition of C5a and C3a has been reported to increase TLR4-induced IL-6 and TNF secretion [49]. Rather than showing direct interaction between complement proteins and TLR4 signaling components, these studies highlight synergy between the signaling pathways of anaphylatoxin receptors and TLR4. A distinction between intracellular or extracellular functions of complement has traditionally not been made. However, using lung epithelial cells expressing solely cytosolic C3, Kuska *et al.* [18] demonstrated that intracellular C3 is required for proper signal transduction through the TLR-NF-κB pathway and subsequent production of chemokines and cytokines. Their study provides evidence that cytosolic C3 is a direct regulator of proteins involved in pro-inflammatory signaling. Our data support this conclusion, although the specific regulatory role of C3 appears to differ between cell types. Our experiments were done in a THP-1 monoculture with heat-inactivated bovine serum, from which internalization of C3 was not observed in B cells or fibroblasts [7, 50]. Thus, our findings can be attributed to cell-intrinsic C3. Though we have not conclusively proved that the attenuation of IFN-β production is solely exerted by intracellular C3, this is supported by two findings: extracellular C3 or NHS failed to rescue the phenotype in C3 KO THP-1-derived macrophages, and an extracellular-only FB-blocking antibody did not reproduce the effects seen with a cell-permeable FB inhibitor. Together, these results point to an intracellular origin. Future work using cells retaining only cytosolic C3, as done by Kuska *et al.* [18], could further validate this conclusion.

We found that IRF3 only co-precipitated with TBK1 in the absence of C3, both at baseline and following LPS-stimulation. Moreover, C3 co-precipitated with IRF3, but not TBK1, in lysates from WT THP-1 cells expressing EGFP-tagged C3. These findings suggest that C3 reduces IRF3 activity by limiting its interaction with TBK1, thus preventing the full phosphorylation of IRF3 which is required for efficient dimerization and nuclear translocation. Other host proteins known to inhibit IRF3 activity primarily target the phosphorylation or degradation of IRF3, such as the phosphatases PP2A [51] or PPM1B [52], or the ubiquitin ligases RAUL [51] and TRIM family proteins [53, 54]. These IRF3-regulating proteins are essential to fine-tune our response to infection, ensuring that the antiviral system remains primed but inactive in the absence of infection [55]. It is conceivable that cell-intrinsic C3 may play a similar role in modulation of the IRF3 response following stimulation of PRRs. Moreover, other host proteins targeting the interaction between TBK1 and IRF3 have been identified. For instance, Li *et al.* reported that the host regulator YWHAZ suppresses the expression of IFN-β induced by RNA viruses by binding IRF3 and inhibiting its interaction with both TBK1 and the nuclear importer KPNA3 [56].

Further investigations revealed that C3 deficiency increased LPS-induced IRF3 phosphorylation at Ser^396^, but not Ser^386^. This was accompanied by enhanced dimerization and nuclear translocation of IRF3. Although Ser^386^ is considered the most important residue for IRF3 dimerization, mutations of Ser^396^ and other residues interacting with phosphorylated Ser^396^ also reduce dimerization and nuclear translocation of IRF3, in addition to inhibiting IFN-β expression by approximately 50% [57]. The precise phosphorylation kinetics of individual IRF3 residues remain unresolved [40]. One model proposes that Ser^386^ phosphorylation follows Ser^396^ phosphorylation, which relieves IRF3 autoinhibition [58]. In contrast, a study of viral infection found the opposite, that Ser^386^ was phosphorylated before Ser^396^, as the latter were present in dimers whereas the former was not [59]. If Ser^396^ phosphorylation represents a later event that requires more sustained engagement with TBK1, this could explain why C3 selectively impairs Ser^396^ phosphorylation: by destabilizing the IRF3–TBK1 complex, C3 may preferentially disrupt the prolonged interaction needed for secondary-site phosphorylation while leaving earlier, more transient phosphorylation events comparatively intact.

A limitation of this work is that most of our mechanistic findings rely on C3 KO clones generated by CRISPR/Cas9, which carries an inherent risk of off-target effects [60]. However, several pathways differentially regulated in our C3 KO THP-1 cells have also been reported in other studies using C3 KOs [18], and key findings were confirmed in C3 deficient BMDMs and in hMDMs, reducing the likelihood of clone-specific artifacts. Moreover, our findings are based on in vitro differentiated macrophages, which may not fully recapitulate complement biology in vivo.

Nonetheless, our findings point to a possible therapeutic avenue. We found that small molecule inhibitors of FD and FB attenuated the production of IFN-β in both primary hMDMs and THP-1-derived macrophages by limiting LPS-induced cleavage of C3. This could be utilized to treat conditions where excessive type I IFNs are considered detrimental. Type I IFNs are largely protective in viral infections, but their role in bacterial infections is less clear. In TB, type I IFNs and related ISGs are considered detrimental at high levels, while tonic or low-level IFN signaling may be protective [42]. Although intracellular growth of *Mtb* was not affected in our system, 3 days post-infection might have been too early to observe effects. Reliable assessment of bacterial burden at later time points was precluded due to cell death and detachment. Additionally, the detrimental role of type I IFNs in *Mtb* infection is likely due to a complex interplay of different immune cells and types of interferons [61]. The suppression of IFN-β by cell-intrinsic C3 could still be exploited as adjunctive host-directed therapy to decrease type I IFN-induced pathophysiology. While there are no published or ongoing clinical trials that target the IFN pathway directly in TB patients, treatment with Janus kinase (JAK) inhibitors has been proposed as a possible host-directed therapy for TB and was found to shorten treatment duration in immunocompetent mice [62]. JAK-inhibitors limit type I IFNs, but also several other JAK/STAT-signaling cytokines that are key in TB, such as IFNg, IL-6, and IL-12 [43]. Although successful treatment for various conditions, JAK inhibitors have been linked to adverse events [63] and infection exacerbation [64]. As C3 plays a wide role in immunity [65], care must be taken to avoid similar effects from prolonged blockage of C3 cleavage. Complement inhibitors can increase the risk of infections [66], but the small-molecule FD inhibitor danicopan used in this work was not found to be associated with such a risk in a clinical trial on patients with paroxysmal nocturnal hemoglobinuria [67].

Dysregulation of type I IFNs is also considered an important factor in the pathogenesis of several autoimmune diseases. For instance, in patients with systemic lupus erythematosus (SLE), increased serum type I IFN activity is associated with disease severity [68], and increased expression of ISGs is frequently observed in PBMCs from patients with active disease [69]. The complement system also plays an important role in the pathophysiology of SLE [70]. C3 serum levels are often used as a biomarker of disease activity in SLE patients, and a recent longitudinal study reported that SLE patients with high type I IFN levels had overall lower C3 serum levels than patients with low type I IFN levels [71]. Although that study measured systemic rather than cell-intrinsic C3, and the mechanistic relationship remains unclear, investigations into the activity of cell-intrinsic C3 in PBMCs from patients with active SLE could be an interesting focus of future studies.

## Materials and methods

### Primary cells

Human buffy coats and serum were obtained from the blood bank at St. Olavs Hospital (Trondheim, Norway). Buffy coats were made from blood samples drawn from healthy volunteers who signed consent for experimental procedures, approved by the Regional Ethical Committee in Central Norway (#S-04114). Human peripheral blood mononuclear cells (PBMCs) were isolated from buffy coats by density gradient centrifugation using Lymphoprep^TM^ (Serumwerk, 1858) as previously described [72]. Monocytes were selected by adherence and then differentiated into human monocyte-derived macrophages (hMDMs) in RPMI 1640 (Sigma-Aldrich, R8758) supplemented with 25 ng/mL M-CSF (R&D Systems, 18340318), 0.68 µM L-glutamine (Sigma-Aldrich, G7513) and 10% pooled human serum. 12 h prior to the start of experiments, the culture media of hMDMs was changed to RPMI 1640 supplemented with 10% heat-inactivated human serum to prevent uptake of exogenous complement components.

### Cell lines

CRISPR-Cas9-gene edited THP-1 cells were purchased from Cyagen. The C3 KO cells have a 17bp deletion in exon 3, while the C3aR1 KO cells have a 2bp deletion in exon 2, and the C5 KO cells have a 1bp insertion in exon 12. All KO cell lines are homozygous single-cell clones, and deletions and insertions were confirmed by Sanger sequencing. Single-cell clones in which the gene-editing was unsuccessful were used as control cells. In experiments without KO cells, THP-1 WT cells (ATCC, TIB-202) were used.

THP-1 cells were cultured in RPMI 1640 supplemented with additional 0.68 µM L-glutamine, 10 mM HEPES (Gibco, 15630-056), 50 µM b-Mercaptoethanol (Gibco, 31350-010), and 10% heat-inactivated fetal bovine serum (FBS, Gibco, 10270) at 37°C and 5% CO2. The cells were maintained at a concentration between 200 000 and 1 000 000 cells/mL. Prior to experiments, the THP-1 cells were differentiated into macrophages using 60 ng/mL phorbol 12-myristate 13-acetate (PMA, Sigma-Aldrich, P1585) for 24 h, followed by 48 h of rest in PMA-free cell culture medium.

### Generation of THP-1 cells expressing C3-EGFP

A lenti-ORF clone encoding C3 (Origene, RC215069L1) with C-terminal Myc- and DDK-tags was used as a backbone to generate a plasmid expressing EGFP-tagged C3 under the control of a EF1α promoter. The CMV promoter in the original plasmid was replaced by EF1α using XbaI and AsiSI restriction sites. EF1α was PCR amplified using lentiCRISPR v2 (Addgene #52961) and the following primers:

EF1a.XbaI_F 5’ CCGGGCCCGCTCTAGAGGGCAGAGCGCACATCGCCCAC 3’

EF1a.AsiSI_R 5’ CCCATGGCGATCGCCATGGTGGCACCGGTAGCGCTAGCCTGTGTTCTGGCGGCAAACCCG 3’

Subsequently, EGFP was cloned on the C-terminal of this plasmid using NotI and EcoRI restriction sites. EGFP was PCR amplified using pEGFP-c1 and the following primers:

EGFP.NotI_F 5’ CGCGTACGCGGCCGCCCGCCACCATGGTGAGCAAGGGCGAGGAGCTGTT 3’

EGFP.PmeI_R 5’ AGGTCGAGAATTCGAACTACTTGTACAGCTCGTCC 3’

To produce pseudoviral particles, HEK293T cells were co-transfected with the C3-EGFP plasmid and the following packaging plasmids; pMDLg/pRRE (Addgene #12251), pMD2.g (Addgene #12259), and pRSV-rev (Addgene #12253), using GeneJuice Transfection Reagent (Sigma-Aldrich, 70967). 24 h after transfection, the media was changed to normal cell culture media. 48 h after media change, the supernatant was harvested and centrifuged at 500 x g for 10 min.

THP-1 WT cells were transduced by spinoculation with pseudoviral particles at 1200 x g for 90 min in the presence of 8 µg/mL (Biosettia, 1,000X). When the cells had grown sufficiently, they were sorted on their GFP-expression using fluorescence-activated cell sorting (FACS). Briefly, cells were spun down and resuspended in cold Dulbecco′s Phosphate Buffered Saline (PBS, Sigma Aldrich, D8537) at 5-10 million per mL, sorted on a BD FACSAria™ Fusion Flow Cytometer (BD Biosciences), resuspended in RPMI supplemented with 30% FBS, and cultivated as normal.

### Murine bone marrow derived macrophages

Female C57BL/6J mice (referred to as WT) and B6.129S4-C3tm1Crr/J (referred to as C3^-/-^) Jax mice were purchased from Charles River Laboratories at 8 weeks of age. Prior to euthanasia, the mice were housed at the Comparative medicine Core Facility (CoMed), at the Norwegian University of Science and Technology (NTNU). The mouse husbandry was in regulations with guidelines from the Federation of European Laboratory Animal Science Associations (FELASA).

After euthanasia by CO2, both hind legs of the mice were excised and cleaned. The femur and tibia were soaked in 96% ethanol, gently cleaned free of muscle and tissue, separated, and transferred to sterile 0.5 mL Eppendorf tubes with a hole in the bottom made with a 19G needle. These tubes were then placed in sterile 1.5 mL Eppendorf tubes containing RPMI 1640 medium supplemented by 10% FBS, 0.68 µM L-glutamine, and 10 mM HEPES (complete medium). Bone marrow cells were flushed out by centrifugation for 15 s at 10,000 x g. Next, red blood cells were lysed by incubation for a few min in eBioscience™ 1X RBC Lysis Buffer Solution (Invitrogen™, 00-4333-57). The remaining cells were spun down and cultivated at 37°C and 5% CO2 in sterile 92 mm^2^ petri dishes (Sarstedt, 82.1473.001) in complete medium supplemented with penicillin/streptomycin (Merck, P0781) and 10 ng/mL murine M-CSF (Merck, M9170) to differentiate them into bone marrow derived macrophages (BMDMs). One day prior to experiments, the BMDMs were gently scraped out of the dishes, counted and seeded at 400 000 cells per well in 24-well cell culture plates, in complete medium without M-CSF and antibiotics. The remaining cells were split 1:2 before reaching 80% confluence.

### Reagents and cell stimulation

Ultrapure K12 LPS from *E. coli* (InvivoGen, tlrl-eklps) was used to stimulate cells at a concentration of 10 ng/mL. The cell-permeable factor B inhibitor, obtained under an MTA with GlaxoSmithKline, and the small molecule factor D inhibitor danicopan (MedChemExpress, HY-117930) were used at a concentration of 10 mM and were added to hMDMs or THP-1 cells 1 h prior to LPS stimulation. The TBK1-IKKε inhibitor MRT67307 was received as a gift from P. Cohen, University of Dundee, [73], and it was used to pre-treat the cells for 1 h at a concentration of 2 µM, before LPS stimulation. DMSO was used as a negative vehicle control for the FB and FD inhibitors, and MRT67307. The anti-IFNAR chain 2 antibody, clone MMHAR-2 isotype IgG2a (EMD Millipore, MAB1155) and the IgG2a κ isotype control antibody (BioLegend, 400224) were used at a concentration of 5 mg/mL and were added to cells 30 min before LPS stimulation. Anti-factor B antibody FB28.4.2 was received as a kind gift from Santiago Rodríguez de Córdoba [38] and used to pre-treat cells for 1 h before LPS stimulation at a concentration of 50 mg/mL. As a control antibody, Ultra-LEAF^TM^ purified mouse IgG2b, k isotype control antibody (BioLegend, 401216) was used at the same concentration.

### RNA sequencing and analysis

Total RNA was isolated from THP-1 cells using the QIAzol lysis reagent (Qiagen, 79306) and chloroform extraction. After cell lysis with 500 µL QIAzol, 100 µL of chloroform (Supelco, 1.02445) was added to the samples, and they were centrifuged for 15 min at 16,100 x g for phase separation. The aqueous upper layer containing the RNA was transferred to separate tubes containing 265 µL 70% ethanol for RNA purification using RNeasy Mini spin columns (Qiagen, 74106) and following the manufacturer’s instructions. DNA was digested using RNase-free DNase (Qiagen, 79256). All downstream processing steps and analysis up to gene Ontology enrichment analysis were carried out by the Genomic Core Facility (GCF) at NTNU. The RNA was quality controlled using an Agilent 2100 Bioanalyzer (Agilent), and RNA quantity was detected by Qubit fluorometric quantitation. All samples had a RIN value greater than 10.0.

RNA sequencing libraries were prepared using the Illumina Stranded mRNA Prep (Ligation) Kit (Illumina, 20040534), following the manufacturer’s protocol. 400 ng total RNA was used as the starting material. First, mRNA was isolated from the total RNA using poly-T oligo-attached magnetic beads, followed by random fragmentation at 94°C for 8 min. First-strand cDNA synthesis was performed at 42°C for 15 min using random hexamer oligonucleotides and Actinomycin D, enhancing strand specificity by supporting RNA-dependent synthesis while preventing spurious DNA-dependent synthesis. Next, second-strand cDNA synthesis was conducted at 16°C for 1 h, during which the RNA template was removed, and a complementary strand was synthesized to create blunt-ended, double-stranded cDNA fragments. To achieve strand specificity, deoxyuridine triphosphate (dUTP) was incorporated instead of deoxythymidine triphosphate (P) during this step, quenching the second strand during amplification. Subsequently, 3’ end adenylation was performed at 37°C for 30 min, followed by the ligation of anchors and Illumina dual index adapter oligonucleotides at 30°C for 10 min. Library fragments underwent cleanup using AMPure XP beads (Beckman Coulter, 31723123) and were enriched through 11 cycles of PCR. The final libraries were purified with AMPure XP beads, quantified using the KAPA Library Quantification Kit (Kapa Biosystems, 07960140001), and validated with the Agilent High Sensitivity DNA Kit (Agilent Technologies, 5067-4627) on a 2100 Bioanalyzer (Agilent Technologies). The size of the DNA fragments was determined to range from approximately 200 to 600 bp, with a peak around 348 bp.

The analysis of the RNA sequencing raw data was performed using the R software (R Core Team 2017), Bioconductor [74] packages including DESeq2 [75, 76] and the SARTools package developed at PF2 - Institut Pasteur. Normalization and differential analysis were carried out according to the DESeq2 model and package. GO enrichment analysis was performed on all differentially expressed genes using the online tool ShinyGO 0.85.1 [77]. Subsequent data organization and visualizations were done in R-3.3.2 using the “dplyr” and “ggplot2” packages. The top 30 upregulated or downregulated KEGG pathways for each condition were selected, and a threshold for FDR of < 0.05 was used. For analysis of ISGs, a complete list of ISGs from Yu, J *et al.* (2025) [78] was used. To find differentially expressed genes for each condition, a threshold for adjusted p-value of < 0.05 was used. Upregulated genes were defined as genes with a |log2FoldChange| > 1, downregulated as genes with a |log2FoldChange| < -1. Volcano plots and heatmap were both created using the “ggplot2” package in R-3.3.2.

### ELISA and multiplex cytokine assays

Quantification of CXCL10 and IFN-β in cell culture supernatants was performed using the following ELISA kits: CXCL10/IP-10 DuoSet ELISA (R&D Systems, DY266-05) and IFNβ DuoSet ELISA (R&D Systems, DY814-05). 10% BSA (Sigma Aldrich, A7030) was used to make reagent diluent (1% BSA in PBS), which was also used for blocking. The ELISA kits were used according to manufacturer’s instructions, except all incubation volumes were reduced to 40 mL, and 100 mL for the blocking step. All ELISAs were performed in half-area 96 well, high binding plates (Corning, CLS3690-100E). 3,3’,5,5’-tetrametylbenzidin (TMB, BioLegend, 421101) was used as a HRP substrate. All plates were washed with a Hydraspeed platewasher (Tecan, 30190101), with an extra washing step in a biosafety cabinet using the same washing buffer when bacteria were present in the supernatant. The iMark microplate reader (Bio-Rad Laboratories, 168-1130) was used to read absorbance at 450 nm with a background read at 570 nm.

Cell culture supernatant from THP-1 cells was analyzed for 27 cytokines using Bio-Plex Pro Human Cytokine 27-plex assay (Bio-Rad Laboratories, M500KCAF0Y). Cell culture supernatant from BMDMs was analyzed with a custom 16-plex ProcartaPlex multiplex (Thermo Fisher, PPX-16-MXZTE3K). Both were done in single replicates using Luminex xMAP Technology on a Bio-Plex 200 System (Bio-Rad Laboratories, 171000201). The manufacturer’s protocol was followed with the recommended concentration of reagents and supernatants, but in reduced volume (1:2). This modification has been used in previous studies without reducing assay performance [79].

### RT-qPCR

Total RNA was isolated from THP-1 cells or hMDMs using QIAzol and chloroform extraction as described under RNA sequencing and analysis. RNA concentrations were measured using NanoDrop 1000 Spectrophotometer (Thermo Fisher Scientific), and then cDNA was produced from total RNA using the High-Capacity RNA-to-cDNA^TM^ kit (Applied Biosystems^TM^, 4388950). Quantitative real-time PCR (qPCR) was performed on 2 ng/mL cDNA with TaqMan™ Fast Advanced Master Mix (Applied Biosystems^TM^, 4444964) in duplicates in a StepOnePlus™ Real-Time PCR cycler (Applied Biosystems^TM^). The relative expression of the genes tested was calculated using the ΔΔCT method, wherein the level of mRNA encoding TATA-box binding protein (TBP) was used as an endogenous control for normalization of results. The following TaqMan Gene Expression Assays (Applied Biosystems^TM^, 4331182) were used: *IFNB1* (Hs01077958_s1), *IRF3* (Hs01547283_m1), *TLR4* (Hs00152939_m1), *TBP* (Hs00427620_m1).

### Immunoblotting

Cell lysates for immunoblotting were prepared by simultaneous extraction of total protein and RNA using the QIAzol lysis reagent and chloroform extraction, or by RIPA lysis buffer. After lysing the cells with 500 µL QIAzol, 100 µL of chloroform was added to the samples, and they were centrifuged for 15 min at 16,100 x g for phase separation. The aqueous upper layer containing the RNA was transferred to separate tubes for RNA purification. 160 µL absolute ethanol was added to the remaining organic phase, which was then centrifuged for 5 min at 2000 x g to pellet the DNA. The supernatant was transferred to tubes containing 750 µL isopropanol (Sigma-Aldrich, 190764) and incubated for 10 min at room temperature, before it was centrifuged for 10 min at 12,600 x g. The protein pellets were then washed three times with 0.3 M guanidine hydrochloride in 96% ethanol, once with absolute ethanol, and then dissolved by heating the pellets to 80 °C for 10 min in a buffer containing 2% SDS and 4 M urea, and NuPAGE LDS Sample Buffer (Invitrogen, NP0008) containing 0.1 M DTT (AppliChem GmbH, A3668,0050). For experiments where RNA was not needed, lysates were made using 1X RIPA lysis buffer (150 mM NaCl, 50 mM Tris-HCL (pH 7.5), 1% Triton X-100, 5 mM EDTA), supplemented with EDTA-free cOmplete™ Mini Protease Inhibitor Cocktail tablets (Roche, 11836170001) and PhosSTOP phosphatase inhibitor cocktail (Roche, 04906837001).

Pre-cast gradient 4-12% Bis-Tris NuPAGE gels (Thermo Fisher, WG1402BOX and NP0322BOX) were used for SDS-PAGE, with either MES (2-100 kDa proteins, Invitrogen, NP000202) or MOPS (100-220 kDa proteins, Invitrogen, NP000102) buffers. Proteins were transferred to nitrocellulose iBlot Transfer Stacks (Invitrogen, IB301031 or IB33002X3) from the gel using the iBlot 2 Gel Transfer Device (Invitrogen, IB21001). The blots were blocked with 5% milk in in Tris-buffered saline-Tween (TBS-T, 20 mM Tris-HCl pH 7.5, 150 mM NaCl, 0.1% Tween-20) for 30 min or 5% bovine serum albumin (BSA) (Sigma-Aldrich, A9418) in TBS-T for 1 h at RT, and then incubated with the primary antibody indicated either overnight or for 48 h at 4°C. The following primary antibodies were used:

| **Antibody** | **Producer** | **Species** | **Catalogue number** | **Dilution** |
| --- | --- | --- | --- | --- |
| C3 | Invitrogen | Rabbit polyclonal | PA5-21349 | 1:1000 |
| C3 | Sigma Aldrich | Goat polyclonal | 204869 | 1:5000 |
| GAPDH | Abcam | Mouse monoclonal | Ab9484 | 1:5000 |
| IRF1 | Cell Signaling<br>Technology | Rabbit monoclonal | 8478 | 1:1000 |
| IRF3 | Abcam | Rabbit monoclonal | Ab68481 | 1:1000 |
| IRF3 | Protein tech | Mouse monoclonal | 16868765 | 1:5000 |
| IRF7 | Cell Signaling<br>Technology | Rabbit monoclonal | 13014 | 1:1000 |
| NF-κB p65 | Cell Signaling<br>Technology | Mouse monoclonal | 6956 | 1:1000 |
| PARP1 | Abcam | Rabbit monoclonal | Ab32138 | 1:2000 |
| Phospho-IRF3 (S386) | Abcam | Rabbit monoclonal | Ab76493 | 1:1000 |
| Phospho-STAT1 (Y701) | Cell Signaling<br>Technology | Rabbit monoclonal | 7649 | 1:1000 |
| Phospho-TBK1 (S172) | Cell Signaling<br>Technology | Rabbit monoclonal | 5483 | 1:1000 |
| Phospho-IRF3 (S396) | Cell Signaling<br>Technology | Rabbit monoclonal | 29047 | 1:1000 |
| STAT1 | BD Biosciences | Mouse monoclonal | 610185 | 1:1000 |
| TBK1 | Cell Signaling<br>Technology | Rabbit monoclonal | 3504 | 1:1000 |
| β-tubulin | Abcam | Rabbit polyclonal | Ab15568 | 1:10 000 |
| β-tubulin | Cell Signaling<br>Technology | Mouse monoclonal | 86298 | 1:5000 |

After incubation with primary antibodies, blots were washed three times for 5 min with TBS-T and then incubated with an appropriate HRP-conjugated secondary antibody for 1 h at RT. The following secondary antibodies were diluted in 1% BSA in TBS-T as indicated: goat anti-mouse (Dako, P044701-2, 1:5000), swine anti-rabbit (Dako, P039901-2, 1:4000), donkey anti-goat (Invitrogen, A16005, 1:5000). Western blots were developed with SuperSignal™ West Femto Maximum Sensitivity Substrate (Invitrogen, 34096) and then visualized using the LI-COR Odessey Fc Imaging System (LI-COR Bio-technology).

### Immunoprecipitation of endogenous proteins

For immunoprecipitations (IPs), a 1X lysis buffer containing 150 mM NaCl, 50 mM Tris-HCl, pH 8.0, 1 mM EDTA, and 0.5% IGEPAL® CA-630 (Sigma-Aldrich, I8896-100ML) was supplemented with EDTA-free Complete Mini protease Inhibitor Cocktail Tablets and a PhosSTOP phosphatase-inhibitor cocktail, as well as 50 mM NaF, and 2 mM Na3VO3. After stimulation, THP-1 cells were washed once with cold PBS and then lysed using the 1X lysis buffer. The samples were incubated in this buffer for 15 min before lysate collection, after which the samples were centrifuged at 16,100 x g for 15 min at 4°C.

Pierce™ Protein Assay Kit (Thermo Scientific™, 23225) was used to normalize the samples based on their total protein level. 90 µg was used for immunoblotting of whole lysates, while 1 mg was used for IPs of endogenous proteins. IPs of endogenous proteins were carried out using 10 µg of antibody per IP sample for α-C3c (Agilent Dako, A006202-2), 2.5 of antibody per IP for α-GFP (Takara, 632381), and 5 µg per IP for α-IRF3 (Abcam, Ab68481), with equal amounts used for the rabbit IgG monoclonal isotype control (Abcam, Ab172730). Invitrogen Dynabeads™ protein A (Thermo Fisher, 10006D) were used to couple rabbit antibodies and Invitrogen Dynabeads™ protein G (Thermo Fisher, 10003D) were used to couple mouse antibodies, together with the corresponding DynaMag™-2 Magnet (Thermo Fisher, 12321D). These were used according to the manufacturer’s instructions, apart from a few steps explicitly stated. The bead incubation time with antibody was increased from 10 to 20 min. After binding of the antibodies to the beads, they were covalently crosslinked using Pierce™ BS^3^ crosslinkers (Thermo Scientific, A39266) at the highest recommended concentration (5mM) with 20mM HEPES with 0.15M NaCl as conjugation buffer. 1M Tris-HCl was used as quenching buffer. The beads were washed twice in the quenching buffer prior to crosslinking to change the buffer system. After quenching, the beads were put in 1X lysis buffer without inhibitors, in which they can be stored for up to 7 days at 4°C. Lysates were then added to beads, and incubated with rotation overnight at 4°C.

After overnight incubation, the co-precipitated complexes from the IPs were washed three times with 1 mL of the ice cold 1X lysis buffer (without inhibitors) and separated from the beads using the DynaMag™-2. Each sample was then eluted in a 1X loading buffer (LDS sample buffer) and incubated 10 min at 80°C. No reducing agent was added to the LDS at this step, to avoid breakage and leakage of the light chain antibodies from the beads. The eluates were transferred to clean tubes and DTT (Sigma-Aldrich, Merck) was added to a 40 mM concentration. Subsequently, the samples were heated for 5 min at 80°C and analyzed by SDS–PAGE and WB. IPs and whole-cell lysate input controls were loaded together onto the gels. MOPS was used as running buffer, and the blots were blocked for 30 min with 5 % milk in TBS-T for α-C3c IPS, and for 1 h with 5% BSA in TBST-T for α-IRF3 IPs.

### Native PAGE

For Native PAGE, a modified version [80] of a previously published protocol was used [81]. Stimulated THP-1 cells were lysed with 1X RIPA lysis buffer (150 mM NaCl, 50 mM Tris-HCL (pH 7.5), 1% Triton X-100, 5 mM EDTA), supplemented with EDTA-free cOmplete™ Mini Protease Inhibitor Cocktail tablets and PhosSTOP phosphatase inhibitor cocktail. Samples were normalized according to their total protein level, measured by Pierce™ BCA Protein Assay Kit. The samples were then diluted 1:1 in 2X Novex™ Tris-Glycine Native Sample Buffer (Invitrogen™, LC2673) supplemented with 1% sodium deoxycholate (Sigma Aldrich, 309-70-100G). 8% Novex Tris-Glycine WedgeWell Gels (Invitrogen, XP00080BOX) were pre-run at 4 °C with 25 mM Tris-192 mM glycine (pH 8.4) containing 0.1% sodium deoxycholate in the cathode chamber for 30 min at 40 mA. Per sample, 15 mg of protein was loaded to the gel and electrophoresed at 4°C for 2 h at 25 mA. Subsequently, the gel was incubated for 30 min at RT in 25 mM Tris-192 mM Glycine (pH 8.4) containing 0.1 % SDS and then transferred to nitrocellulose iBlot Transfer Stacks using the iBlot 2 Gel Transfer Device. The blots were blocked with 5% milk in TBS-T for 30 min at RT, incubated with an anti-IRF3 antibody (Abcam, Ab68481, diluted 1:1000) for 48 h at 4°C, and then developed as described under Immunoblotting. For loading control, the same samples were diluted in LDS sample buffer with DTT and analyzed with SDS-page of IRF3.

### Cellular fractionation

Stimulated THP-1 cells were detached by Accutase (Invitrogen, 00-4555-56) and collected by centrifugation (600 x g, 5 min, 4 °C). The cell pellets were washed once with PBS containing 2% FBS. For preparation of total lysate, 1/3 of each sample was centrifuged (600 x g, 5 min, 4 °C) and then lysed by RIPA lysis buffer (150 mM NaCl, 50 mM Tris-HCl (pH 7.5), 1% Triton-X100. The remaining portion of each sample was centrifuged and then resuspended in buffer A (50 mM NaCl, 10 mM (pH 8), 500 mM sucrose, 1 mM EDTA, 0.2% Triton-X100), supplemented with PhosSTOP phosphatase-inhibitor cocktail and EDTA-free Complete Mini protease Inhibitor Cocktail Tablets. The samples were then vortexed and centrifuged (5000 rpm, 5 min, 4 °C). Cytosolic fractions in the supernatants were transferred to clean Eppendorf tubes, while the nuclear fractions were resuspended in buffer B (50 mM NaCl, 10 mM HEPES (pH 8), 25% glycerol, and 0.1 mM EDTA), and centrifuged (5000 rpm, 5 min, 4 °C). Supernatants were discarded and the pellets were resuspended in buffer C (350 mM NaCl, 10 mM HEPES (pH 8), 25% glycerol, 0.1 mM EDTA, Benzonase endonuclease (Millipore, E1014-25KU), supplemented with PhosSTOP phosphatase-inhibitor cocktail and EDTA-free Complete Mini protease Inhibitor Cocktail Tablets). Cytosolic and nuclear fractions were then centrifuged at 15,000 rpm for 15 min to extract proteins by transferring the supernatant to clean Eppendorf tubes.

### Cultivation of and infection with *E. coli* DH5a

Live (DH5a*) E. coli* expressing GFP pZE27GFP (Addgene plasmid 75452) was a gift from James Collins [82]. It was cultivated in Luria-Bertani (LB) medium containing 10 g/L Oxoid™ tryptone (Thermo Fischer, LP0042B), 5 g/L Oxoid™ Yeast Extract Powder (Thermo Fischer, LP0021B), and 10 g/L NaCl. On the day of the experiment, overnight cultures of DH5a were diluted 1:30 in LB medium and incubated at 37°C, 250 rpm to obtain an optical density at 600 nm (OD600) of 0.3-0.4. The bacterial cultures were then centrifuged at 4754 x g for 15 min and washed twice with PBS, after which they were resuspended in RPMI supplemented with either 1 or 10% FBS. The OD600 was measured again after washing, and the bacteria were added to THP-1 cells at a multiplicity of infection (MOI) of 10.

### Cultivation of and infection with *Mycobacterium tuberculosis*

The double auxotrophic *Mtb* H37Rv mc^2^6206 strain (H37Rv ΔpanCD ΔleuCD) was provided by William Jacobs at the Albert Einstein College of Medicine [83, 84]. It was cultivated at 37°C in Middlebrook 7H9 medium made with Middlebrook 7H9 Broth Base (Sigma-Aldrich, M0178) supplemented with 10% Middlebrook OADC (BD Difco, 211886), 0.2% glycerol (Sigma Aldrich, G5516), 0.05% Tween-80 (Sigma Aldrich, P4780), 50 µg/mL L-leucine (Sigma Aldrich, L-8912-25G) and 24 µg/mL D-pantothenate (Sigma Aldrich, P5155-100G). 0.1 mM Sodium propionate (Sigma-Aldrich, 18108) was added to sustain PDIM levels [85]. *Mtb* mc^2^6206 was transformed to express firefly luciferase under the control of a synthetic promoter (MOP) using the pMH109 plasmid (kindly provided by Mark Hickey/David Sherman, Seattle, USA) [86]. The bacterial cultures were kept in log phase (OD600 lower than 0.8) at 37°C with shaking (250 rpm) [86]. The bacterial cultures were kept in log phase (OD600 lower than 0.8) at 37°C with shaking (250 rpm).

For experiments, *Mtb* H37Rv mc^2^6206 with a luciferase construct was cultivated to an OD600 of between 0.2 and 0.6 before the required number of bacteria were spun down for 5 min at 4754 x g. The bacteria were opsonized in sterile-filtered pooled human A^+^ serum for 10 min, washed with 5 mL of PBS, pelleted by centrifugation at 4754 x g for 5 min, and resuspended in RPMI with 10% FBS. For declumping, the bacterial suspension was sonicated for 1 min at power setting 9 using a VWR USC 1200 THD sonicator (Avantor), followed by vortexing for 3×5 s. Bacterial aggregates were subsequently removed by centrifugation at 300 x g for 4 min. OD600 was then measured, and the bacterial suspension was diluted to the desired concentration. An MOI of 1 was assumed to correspond to 1×10^8^ bacteria per mL. Macrophages were infected with *Mtb* at an MOI of 10 for indicated durations at 37°C and 5% CO2. For growth assay experiments, extracellular bacteria were then removed by medium replacement, and the infection was allowed to proceed for the indicated durations.

### Mycobacterium tuberculosis luciferase growth assay

*Mtb* firefly luciferase activity was measured as a proxy to assess the bacterial burden at given time points. Passive Lysis 5X Buffer (Promega, E1941) was diluted to 1X directly in the wells at the given time points. No extra washing step was performed to avoid detachment of cells. After a few min of incubation with the lysis buffer, the cells were scraped and vigorously pipetted to ensure full lysis. 50 µL of sample was then mixed with 50 µL of in-house luciferase assay reagent in a white opaque 96w plate (Thermo Scientific™, 236108). After incubating for 7 min, the luminescence was read on Infinite M Plex (Tecan) with the Tecan Magellan Pro software (version 7.5, Tecan, 2022) and an integration time of 1000 ms.

Luciferase assay reagent was made by mixing 20 mM Tricine, 2.67 mM MgSO47H2O, 0.1 mM EDTA, 33.3 mM DTT, 530 µM ATP and 270 µM Acetyl CoEnzyme A (Lithium salt, Sigma, A2181). dH2O was added to make volume up to 222.6 mL. Finally, 570 mL of 2M NaOH and 1.21 mL of 50 mM Magnesium Carbonate Hydroxide (MgCO3)4Mg(OH)25H2O were added.

### LDH assay

Following infection with *Mtb* or *E. coli*, the cell culture supernatant was collected, and LDH release was quantified using CyQUANT™ LDH Cytotoxicity Assay (Invitrogen, C20300) and following the manufacturer’s instructions. The iMark microplate reader was used to read absorbance at 490 nm with a background read at 655 nm. Cytotoxicity was calculated as the percentage of LDH release compared to a maximum LDH activity control, in which the cells were lysed with 10X Lysis Buffer from the assay kit. In experiments where LDH release was measured, serum levels were reduced to 1% to limit the background signal caused by endogenous LDH activity present in serum, unless cells were cultured for 24h.

### Statistical analyses and graphs

All data analysis and plotting of graphs were performed using Prism (GraphPad, v. 11). Either two-way ANOVA with multiple comparisons (Dunnett’s, Šídák’s or Tukey’s multiple comparisons test) or unpaired t-tests were used for statistical analyses, where the specifics are described in the figure legends. Statistical significance was defined as * = p < 0.05, ** = p < 0.01, *** = p < 0.001, **** = p < 0.0001. Data were expressed as mean ± SEM, and each n represents a biological replicate.

## Supplementary materials

Figures S1 to S4

## Acknowledgments

RNA library prep, sequencing and parts of the bioinformatics analysis were performed in close collaboration with the Genomics Core Facility (GCF), Norwegian University of Science and Technology (NTNU). GCF is funded by the Faculty of Medicine and Health Sciences at NTNU and the Central Norway Regional Health Authority. Housing and euthanasia of mice were provided by the Comparative medicine Core Facility (CoMed), NTNU. CoMed is funded by the Faculty of Medicine at NTNU and Central Norway Regional Health Authority. The factor B antibody was provided by Prof. Santiago Rodriguez de Cordoba (Centro de Investigaciones Biologicas Margarita Salas), while the cell permeable factor B inhibitor was provided by Dr. Claudia Kemper (National Institutes of Health, National Heart, Lung, and Blood Institute).

## Author contributions

THF, TE and HH conceptualized and supervised the project and acquired funding. SK, CÅ, MY, LR, IF, KR, and SU performed experiments and analyzed the data. KR and CK provided resources and discussions of results. SK and CÅ prepared the figures and wrote the original manuscript draft; THF, TE, MY and HH reviewed and edited the manuscript. All authors reviewed and approved of the final manuscript.

## Data, code, and materials availability

Source data including unprocessed gels and blots, statistics, raw data and quantifications used in plotted graphs will be provided with the published manuscript in an Excel Workbook. RNA sequencing data will be deposited in a repository. The code used for analysis will be made available. The factor B antibody was provided by Prof. Santiago Rodriguez de Cordoba, while the cell permeable factor B inhibitor was provided by Dr. Claudia Kemper under an MTA with GSK.

## Supplementary Figures

**Figure S1:**
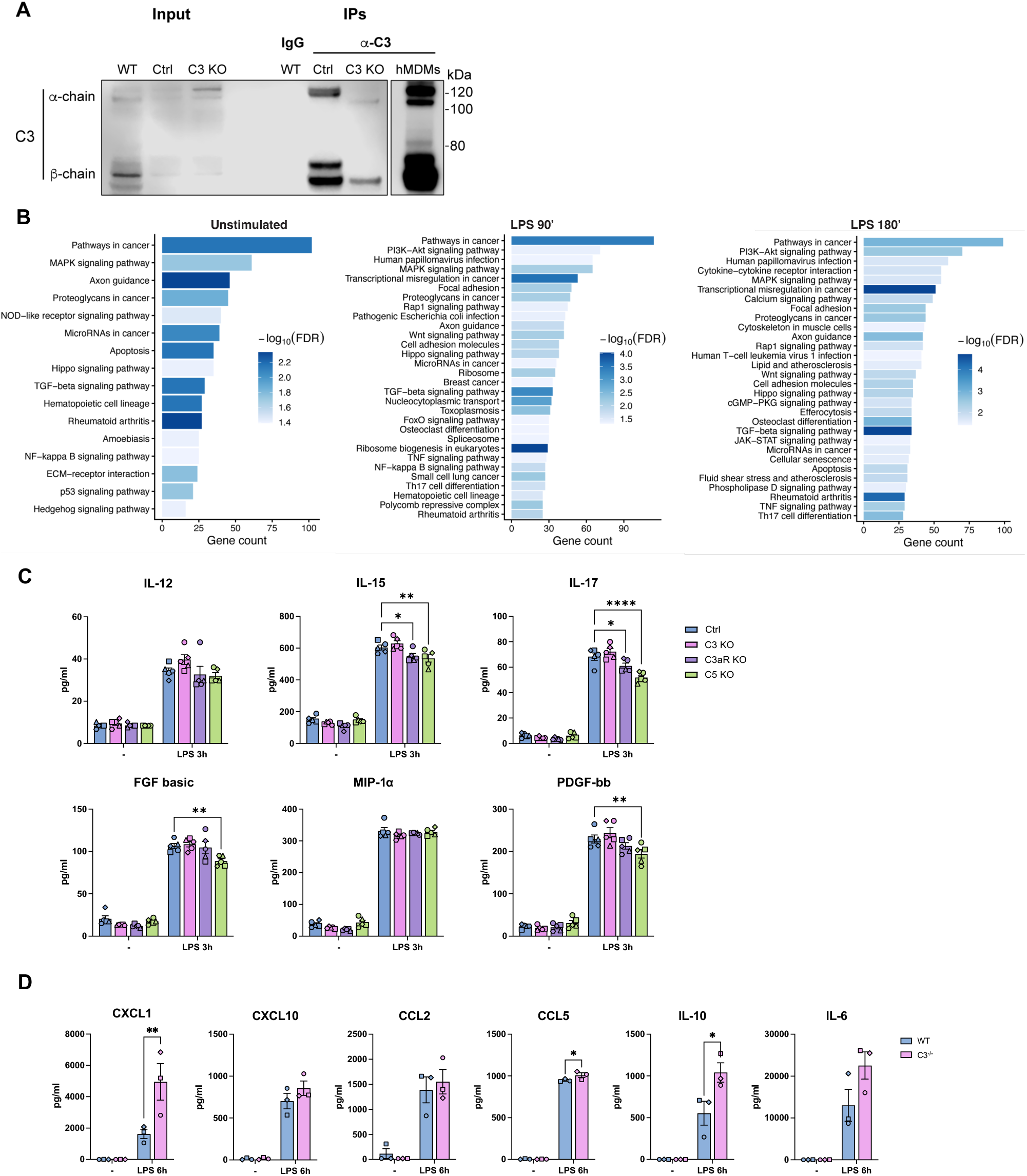
Additional characterization of C3, C3aR and C5 deficient THP-1 cells and C3^-/-^ BMDMs. A) Verification of C3 deficiency in the C3 KO THP-1-derived macrophages by immunoprecipitation of C3. n = 1. B) Gene ontology (GO) pathway analysis of downregulated (adjusted p-value < 0.05, |log2FC| > 2) C3 KO cells compared to control cells, showing the top 30 downregulated pathways in unstimulated cells and cells stimulated for 90 or 180 min with LPS. For the unstimulated condition, only 16 pathways were significantly downregulated. n = 5. C) Cytokines and chemokines which were not significantly increased or decreased in C3 KO THP-1-derived macrophages compared to control. n = 5. D) Quantification of protein levels of CXCL10, CCL2, CCL5, IL-10 and IL-6 in supernatants from BMDMs from WT and C3^-/-^ C57BL/6 mice after 6 h of LPS stimulation. Measured by multiplex immunoassay. n = 3. Data are expressed as mean ± SEM, n denotes biological replicates. Statistical significance was determined using two-way ANOVA and Dunnett’s multiple comparisons test (C) or Šídák’s multiple comparisons test (D). * = p < 0.05, ** = p < 0.01, *** = p < 0.001, **** = p < 0.0001.

**Figure S2:**
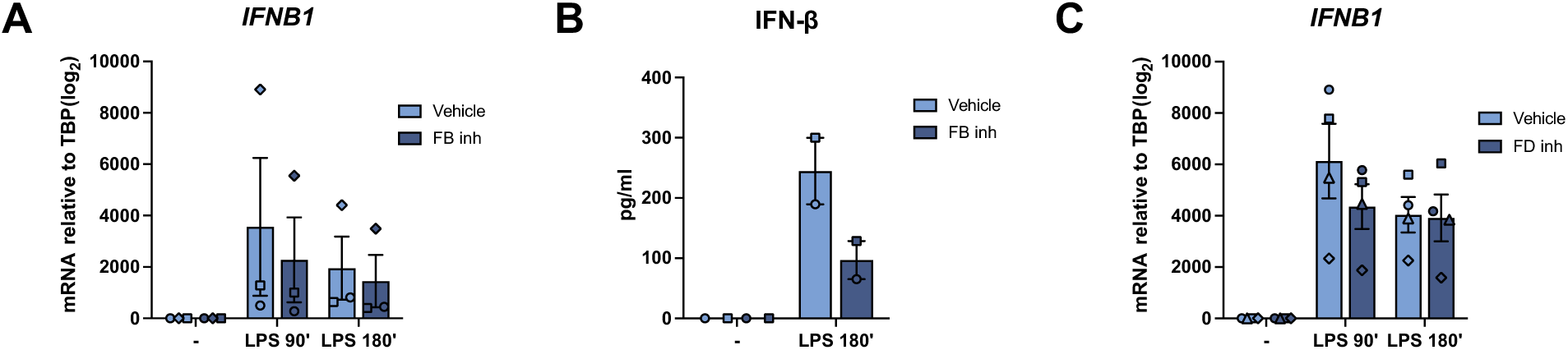
Additional non-normalized quantifications of *IFNB1* and IFN-β levels in hMDMs. A) Quantification of mRNA levels of *IFNB1* in hMDMs by RT-qPCR after 1h of pre-incubation with FB inhibitor and 90 or 180 min LPS stimulation. n = 3. B) Quantification of protein levels of IFN-β in supernatants from hMDMs after 1h of pre-treatment with FB inhibitor and 180 min of LPS stimulation. n = 2. C) Quantification of mRNA levels of *IFNB1* in hMDMs by RT-qPCR after 1h of pre-incubation with FD inhibitor and 90 or 180 min of LPS stimulation. n = 4. Data are expressed as mean ± SEM, n denotes biological replicates. Statistical significance was determined using two-way ANOVA and Šídák’s multiple comparisons test. * = p < 0.05, ** = p < 0.01, *** = p < 0.001, **** = p < 0.0001.

**Figure S3:**
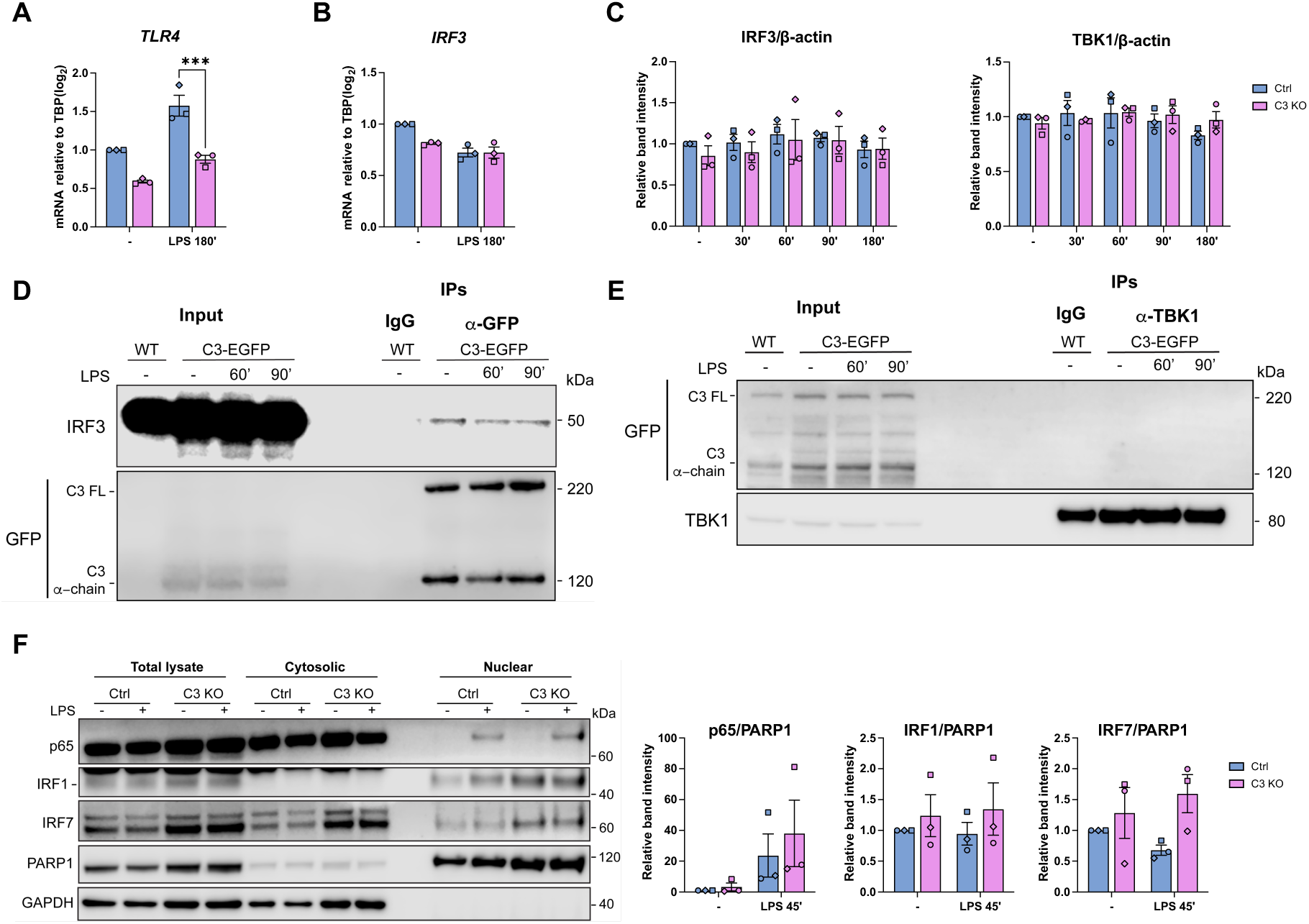
Further characterization of IRF signaling pathways in C3-deficient macrophages. A) Quantification of mRNA levels of *TLR4* by RT-qPCR after 180 min of LPS stimulation in control and C3 KO THP-1-derived macrophages. n = 3. B) Quantification of mRNA levels of *IRF3* by RT-qPCR after 180 min of LPS stimulation in control and C3 KO THP-1-derived macrophages. n = 3. C) Quantifications of total TBK1 and IRF3 protein levels normalised to b-actin protein levels from after LPS stimulation as indicated in control and C3 KO THP-1-derived macrophages, based on densitometric analysis of blots in Figure 4C. n = 3. D) Co-IP of GFP and IRF3 from THP-1 WT-derived macrophages expressing EGFP-tagged C3, after stimulation with LPS as indicated. Quantifications are based on densitometric analysis, normalised to the GFP IP. n = 2. E) Co-IP of TBK1 and GFP from WT THP-1-derived macrophages expressing EGFP-tagged C3, after stimulation with LPS as indicated. n = 1. F) Immunoblot of total lysate, cytosolic fractions and nuclear fractions from control and C3 KO THP-1-derived macrophages stimulated with LPS for 45 min. n = 3. Quantifications are based on densitometric analysis and p65, IRF1 and IRF7 protein levels in the nucleus are normalised to the corresponding nuclear PARP1 band intensity. n = 3. Data are expressed as mean ± SEM, n denotes biological replicates. Statistical significance was determined using two-way ANOVA and Šídák’s multiple comparisons test. * = p < 0.05, ** = p < 0.01, *** = p < 0.001, **** = p < 0.0001.

**Figure S4:**
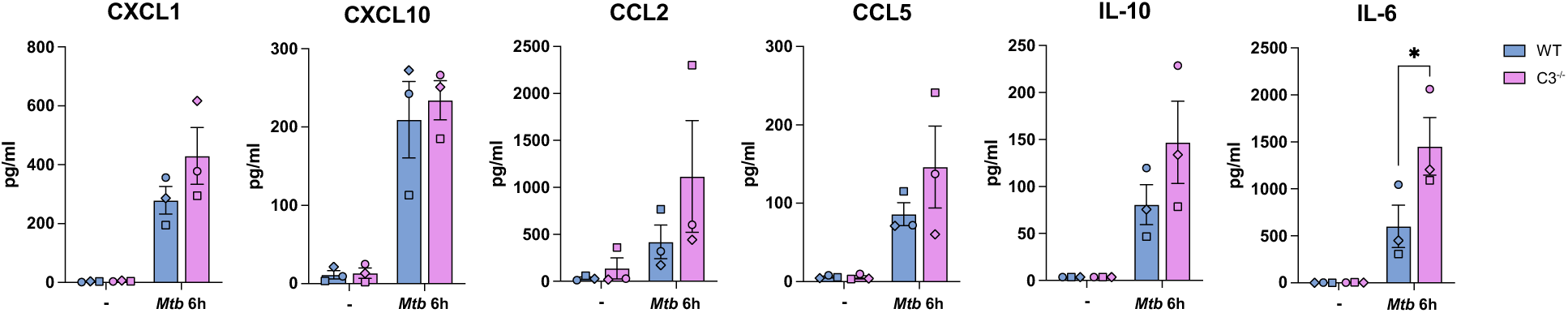
Additional cytokines from multiplex of supernatant from *Mtb*-infected C3^-/-^ BMDMs. Quantification of protein levels of CXCL1, CXCL10, CCL2, CCL5, IL-10, and IL-6 in supernatants from BMDMs from WT and C3^-/-^ C57BL/6 mice after infection with auxotroph *Mtb* for 6 h. Measured by multiplex immunoassay. n = 3 biological replicates. Data are expressed as mean ± SEM. Statistical significance was determined using two-way ANOVA and Šídák’s multiple comparisons test. * = p < 0.05, ** = p < 0.01, *** = p < 0.001, **** = p < 0.0001.

